# This is SPRTA: assessing phylogenetic confidence at pandemic scales

**DOI:** 10.1101/2024.10.21.619398

**Authors:** Nicola De Maio, Nhan Ly-Trong, Bui Quang Minh, Nick Goldman

**Affiliations:** European Molecular Biology Laboratory, European Bioinformatics Institute, Wellcome Genome Campus, Hinxton, Cambridgeshire, CB10 1SD, UK; School of Computing, College of Engineering, Computing and Cybernetics, Australian National University, Canberra, ACT 2600, Australia

## Abstract

Phylogenetics plays a central role in evolutionary biology and genomic epidemiology. Assessing phylogenetic confidence and reliability is therefore crucial and methods to do this, such as Felsenstein’s bootstrap, are among the most used in modern science. However, methods based on Felsenstein’s bootstrap suffer from excessive computational demand, and are unsuitable for large datasets. Furthermore, most of these methods emerge from a cladistic framework which makes their results hard to interpret in the context of genomic epidemiology.

We propose SPRTA (“ SPR-based Tree Assessment”), an efficient and interpretable approach to assess confidence in phylogenetic trees. SPRTA shifts the paradigm of phylogenetic support measurement from evaluating the confidence in clades (groupings of taxa) to genome evolution histories, for example assessing if a lineage evolved from another considered lineage or not. This focus on evolutionary histories is particularly valuable in genomic epidemiology, where typically the evolutionary and transmission history of a pathogen are of interest, not clade content.

We illustrate the use of SPRTA by investigating a global SARS-CoV-2 phylogenetic tree relating *>* 2M genomes, highlighting plausible alternative evolutionary origins of many SARS-CoV-2 variants. We have implemented SPRTA within the free and open source maximum likelihood phylogenetic software MAPLE, available from https://github.com/NicolaDM/MAPLE.

## 1 Introduction

Analysis of genomic data often relies on phylogenetics. Most phylogenetic methods scalable to large datasets, such as maximum likelihood, parsimony-based and heuristic approaches, typically estimate a single phylogenetic tree without intrinsically assessing inference reliability or uncertainty. This issue is typically addressed with methods like Felsenstein’s bootstrap[1] (“ FB”), which has consequently become one of the most cited works in science[2]. For a given dataset, FB typically creates between 100 and 1000 replicates by randomly resampling alignment columns with replacement; phylogenetic inference is performed on each, to estimate replicate trees. The support score of a clade (the group of taxa that are inferred to be all the descendants of one ancestor in the tree) is defined as the proportion of replicate trees containing that clade.

FB, like most well-established phylogenetic methods, has been developed in the context of inter-species evolutionary biology. Consequently, FB has a number of drawbacks when applied to genomic epidemiological datasets:

- Repeatedly performing phylogenetic estimation on all replicate datasets can be excessively computationally demanding. For this reason a number of FB approximations have been proposed in recent years[3–5]; however, these are still not feasibly applicable at pandemic scales (datasets of millions of genomes); see below.
- “ Rogue taxa”, that is, sequences with high phylogenetic uncertainty such as partial sequences or recombinants, can substantially lower the FB support of internal branches of phylogenetic trees[6].
- FB does not measure posterior probability, but repeatability[7]. In genomic epidemiology, phylogenetic branch lengths are usually short, and a single mutation is typically sufficient to define a clade with negligible uncertainty. However, in this scenario FB usually requires 3 mutations supporting any one clade to be able to assign 95% support to it (i.e. so that at least one such mutation will be sampled in *>* 95% of bootstrap alignments), making it excessively conservative in this scenario[1, 8]. Bayesian phylogenetic methods[9, 10] resolve this problem, but are too computationally demanding for large datasets.
- FB, and most other branch support measures, have a “ cladistic” focus, in that they aim at assessing clade inference. While clades have a clear and important interpretation in taxonomy, they are not as relevant in genomic epidemiology, where the focus of phylogenetic inference is typically the reconstruction of the mutation and transmission history of the pathogen, and the assignment of samples to lineages[11, 12].

Existing local branch support measures[13–16] are considerably more computationally efficient than FB, but also rely on a cladistic interpretation. They usually compare the likelihood of the inferred phylogenetic tree against the likelihoods of similar alternative trees. If slightly altering the topology of a maximum likelihood tree, specifically by substituting a clade under consideration, leads to only a small likelihood score reduction, then local branch support measures give a low support score to that clade, meaning that it is considered non-reliable. Local branch support measures are particularly appealing because of their computational efficiency (usually requiring less resources than inference of the phylogenetic tree itself) and because of their robustness to rogue taxa.

Here, we present a new measure of branch support, SPRTA (“ SPR-based Tree Assessment”), that shifts the focus from a “ cladistic” point of view to one centred around mutational histories (“ mutational” focus). We draw heavily on concepts from local branch support measures, and particularly aBayes[14], and as such our approach is inherently robust to rogue taxa. Differently from aBayes, however, we use subtree prune and regraft (SPR; see [17]) moves instead of nearest neighbour interchanges (NNI; [17]). By implementing SPRTA within the pandemic-scale phylogenetic software MAPLE[18] (https://github.com/NicolaDM/MAPLE) we can robustly and interpretably assess phylogenetic reliability at pandemic scales.

## 2 Results

### 2.1 Principles of SPRTA

Given a multiple sequence alignment *D* and a phylogenetic tree *T* inferred from *D*, such as a maximum likelihood pandemic-scale rooted tree inferred with MAPLE[18], our aim is to assign confidence scores to branches *b* of *T*, assessing the reliability of the mutation events inferred along branch *b*. More specifically, we want to assess the reliability of the statement that the genome *B* at the lower end of *b* (the root genome of *S*_*b*_, corresponding to the bottom end of the red branch in Fig. 1) evolved from the ancestor genome *A* at the upper end of *b* (corresponding to the top end of the red branch in Fig. 1). In other words, we want to assesses the probability that indeed *B* evolved from *A*. We call this a “ mutational” branch support measure, as opposed to the “ cladistic” focus on the content of *S*_*b*_ of traditional branch support measures.

**Figure 1:**
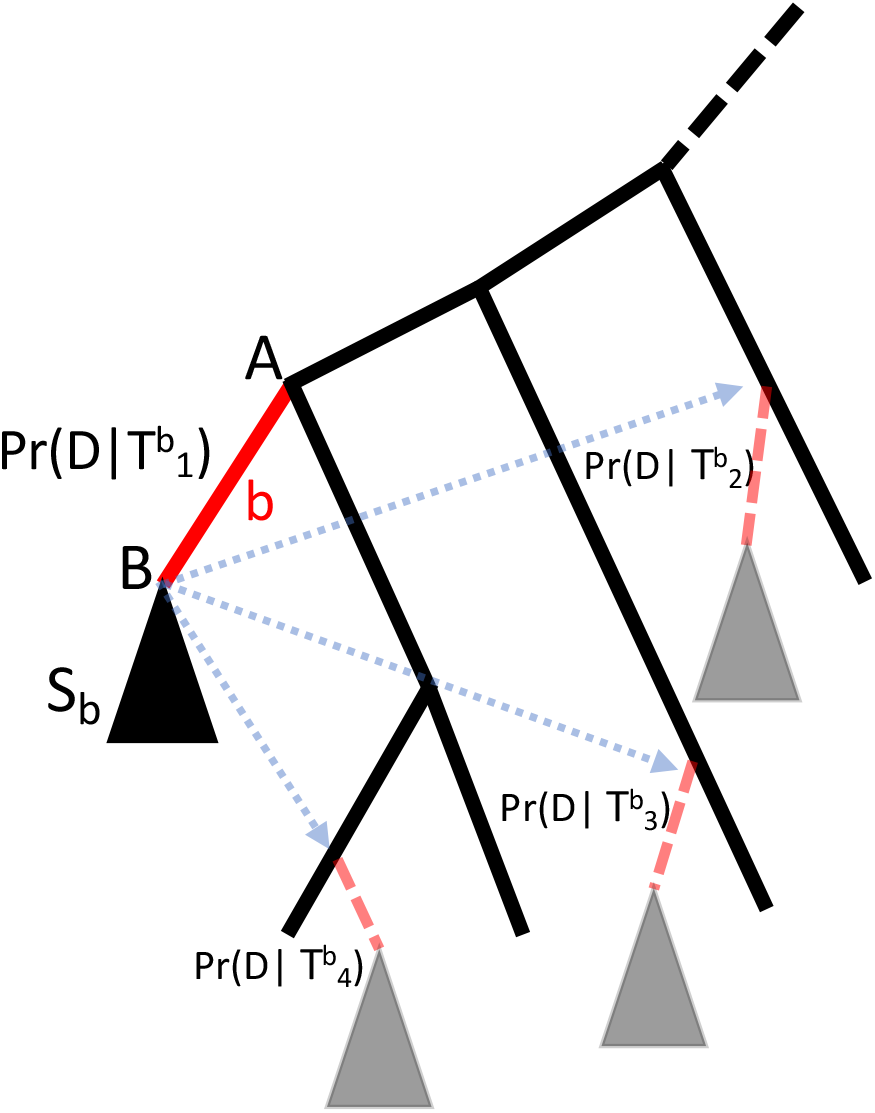
Graphical representation of SPRTA branch support measurement. Here we represent SPRTA branch support calculation for an example tree branch *b* (highlighted as a solid red line). The subtree *S*_*b*_ below *b* is represented with a black triangle and is not affected by any of the SPR moves considered to evaluate the support of *b*. A and B represent the genomes separated by branch *b*. Solid black lines represent branches of the original tree 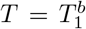 being assessed, while the dashed black line represents the rest of *T* (which is an arbitrarily large tree), not shown here. Possible SPR moves are highlighted with dotted blue arrows, and cause hypothetical new branches (shown as shaded red dashed lines), leading to eventual copies of *S*_*b*_ (grey triangles). The relative likelihood of the original tree is 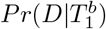 while the likelihoods of the alternative topologies are represented by 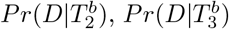, and 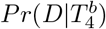. In this case, the support of branch *b* will be 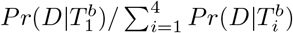. In practice, a large number of alternative topologies might be considered for a starting branch *b* (see Section 4.1).

Given branch *b*, and subtree *S*_*b*_ containing all descendants of *b*, SPRTA considers a number *I*_*b*_ of alternative topologies 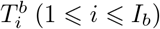 of *T* obtained by performing single SPR moves that relocate *S*_*b*_ as a descendant of other parts of *T* not in *S*_*b*_ (Fig. 1). (See Section 4.1 for the definition of *I*_*b*_.) For notational convenience we assume that 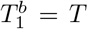 is the original tree topology. The likelihood *Pr*(*D*|*T*^*b*^) of such topologies is routinely and efficiently calculated by MAPLE (up to a constant common multiplicative factor) and is used by SPRTA to define the support score *Pr*(*T* |*D*)_*b*_ of *b*:

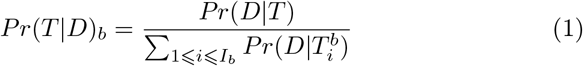

A more detailed description of SPRTA is given in Section 4.1

SPRTA mutational branch support scores need to be interpreted very differently than those of existing, cladistic branch support methods. SPR moves involving branch *b* preserve the clade defined by *b* (Fig. 1), and so SPRTA scores do not represent an assessment of the clade defined by *b*. Rather, SPRTA scores are a particularly useful assessment of tree reliability in the context of genomic epidemiology, where short branches cause low uncertainty in ancestral genomes within a given tree, while the placement of individual incomplete sequences (“ rogue taxa”) can be highly uncertain. SPRTA scores are expected to be robust to rogue taxa, since their placement is expected to have negligible impact on relative likelihood scores and ancestral genome reconstruction at internal tree nodes.

SPRTA scores for the terminal branches of a tree evaluate the placement probability of individual observed genome sequences. In contrast, cladistic support methods do not assess terminal branches and sequence placements.

### 2.2 Computational demand

The SPR search required by SPRTA is typically performed as part of the phylogenetic tree search in many maximum likelihood phylogenetic methods such as RaxML[19] and MAPLE, and so it is not expected to lead to significant additional runtime when executed in conjunction with phylogenetic inference. For comparison, we assessed the computational demand of SPRTA against the main existing measures of branch support (FB[1], LBP[13], aLRT[14], aLRT-SH[15], aBayes[16], TBE[6], and UFBoot2[5]) in measuring the support of branches in trees inferred by MAPLE. SPRTA reduces runtime and memory demands by at least two orders of magnitude compared to all other considered methods, with the difference growing as dataset size increases (Fig. 2). Note also that the premature termination of the lines in Figure 2 for the other methods indicates cases where they could not be run successfully. Among these other methods, those based on FB (FB, UFBoot and TBE) have substantially higher computational demand than local branch support measures.

**Figure 2:**
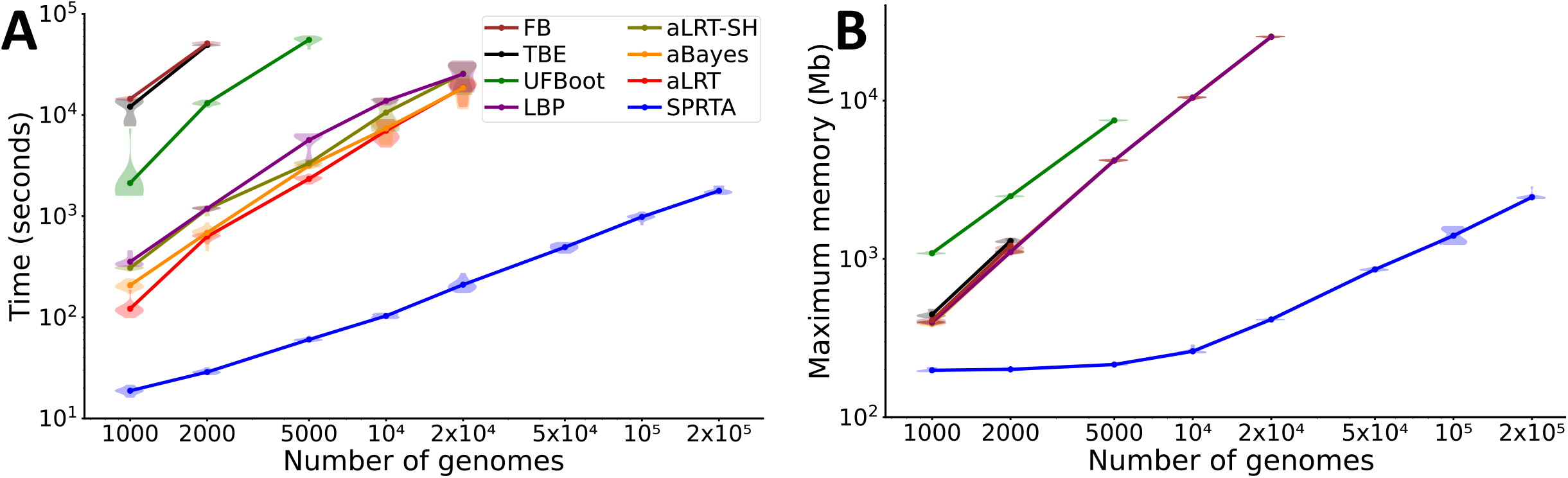
Computational demand. **A** Time and **B** maximum RAM usage of different branch support methods. On the X axis are the number of simulated SARS-CoV-2 genomes (see Section 4.5.2) included in each replicate. For each dataset size considered we ran 20 replicates. Dots show the mean across replicates, while violin plots show variation between replicates. We used the SPRTA implementation within MAPLE v0.6.8, while for all other methods (LBP[13], aBayes[16], aLRT[14], aLRT-SH[15], TBE[6], FB[1], and UFBoot2[5]) we used their implementation within IQ-TREE[20] v2.1.3. For all methods we used the tree estimated by MAPLE as starting tree to reduce computational demand of IQ-TREE branch support methods and to increase comparability between the results of SPRTA and implementations in IQ-TREE. All analyses were run on one core of an Intel Xeon Gold 6252 processor @ 2.10 GHz.

### 2.3 Accuracy

To benchmark different branch support methods, we simulated realistic SARS-CoV-2 genome data (Section 4.5.2) for which we know the true tree and the true mutational history along the tree. Unlike traditional cladistic branch support methods, we do not aim at assessing the reliability of phylogenetic clades. Instead, the focus of our mutational branch support measure is directed at assessing the evolutionary history of the genome. As such, we do not consider simulated clades and topologies when evaluating methods, but only the genome evolutionary history along the true tree. Therefore, comparisons in this section should not be considered as an evaluation of cladistic measures in evaluating clade support (and so they are not an evaluation of these methods *per se*), but as an evaluation of their scores in “ mutational” terms.

In short, we define correctness of phylogenetic tree estimation in terms of the correctness of the genome evolutionary history implied by the inferred phylogenetic tree. Each individual branch of a tree inferred from simulated data is considered correct if the mutations inferred on that branch actually happened in our simulations, and so if the genome at the lower end of the branch actually evolved from the one at the top of the branch. See Section 4.2 for more details on how we define accuracy of branch support. Below, all branch support measures are evaluated based on their ability to assign higher support to correctly inferred mutation events, and lower support to erroneously inferred mutation events.

Generally, all methods give high support scores (*>* 80%) to all mutations, both correctly and wrongly inferred ones (Fig. 3). This is to be expected given that by definition these mutations have been inferred by maximum likelihood, and so will typically have higher likelihood than alternative mutation histories. However, SPRTA is the only method that reliably assigns higher support to correctly inferred mutation events (typically a mean of 98–99%) and lower support scores to wrongly inferred ones (typically a mean of 85–90%): see Figure 3 and Supplementary Figure S1. In fact, while some methods like UDBoot, aLRT, and aBayes give higher support to correctly inferred mutations than does SPRTA, they also similarly support wrongly inferred mutations, and so fail to discriminate between the two. It has to be remembered, however, that cladistic approaches have been developed with a different interpretation of branch support scores, and so these results are not indicative of their performance for that task.

**Figure 3:**
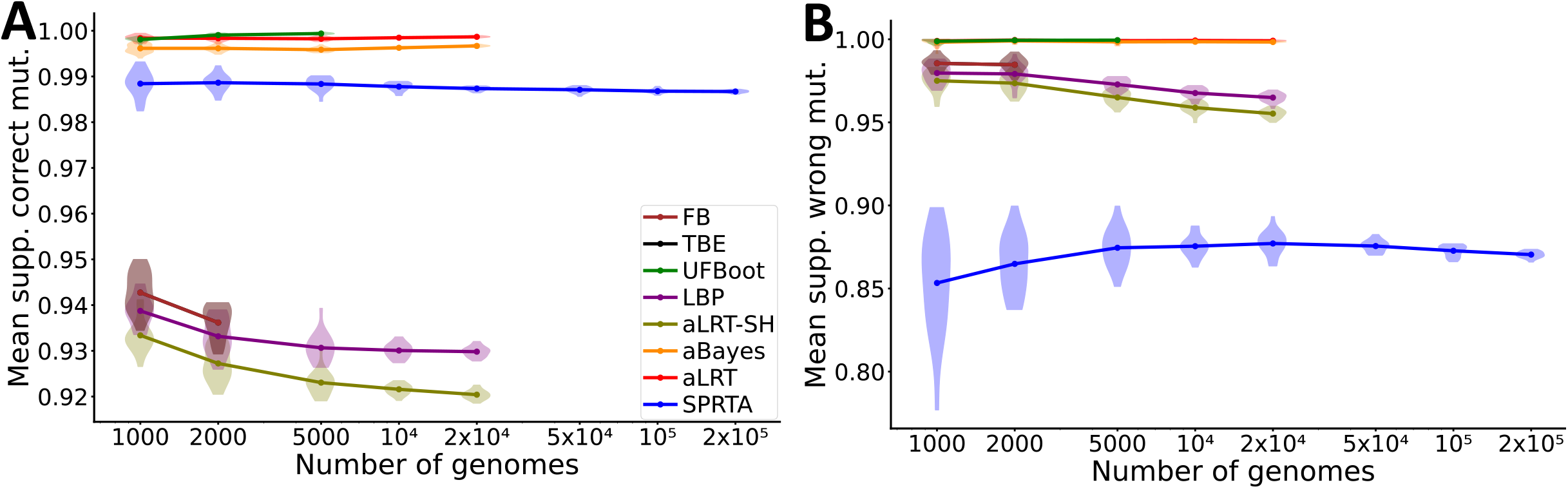
Benchmark of branch support methods. **A** Mean support of correctly inferred mutation events within each replicate. **B** Mean support of wrongly inferred mutation events within each replicate. All other details are as in Figure 2. Branch support scores for methods other than SPRTA were again only calculated when computationally feasible.

### 2.4 Assessing uncertainty in SARS-CoV-2 global evolution

We applied SPRTA to a global public dataset of 2,072,111 SARS-CoV-2 genomes (see Section 4.5.1), too large for the feasible application of any existing cladistic branch support measure. Phylogenetic tree estimation with MAPLE, parallelized over 14 cores of an Intel Xeon Gold 6252 processor @ 2.10 GHz, took around 10 days[21]. Post hoc SPRTA branch support assessment, re-performing SPR move evaluations that had already been performed during tree inference, required 7h:27m on a single core and maximum memory 26.93Gb. This branch support assessment can be also performed simultaneously with tree inference, at negligible additional computational cost compared to the tree inference itself. During tree inference, however, we do not evaluate possible alternative placements of genomes that are less informative than other genomes in the dataset, since alternative placements of these genomes do not increase the likelihood of the considered tree. To evaluate alternative placements of all genomes, for example to identify a larger number of rogue taxa in our dataset, we also performed a more in-depth and computationally demanding SPRTA assessment that did not collapse genomes less informative than other genomes in the dataset; this required 22h:42m on a single core and maximum memory 27.75Gb.

Of the 2,072,111 genomes considered here, 636,022 are mutationally informative (the branch separating them from the tree has inferred mutations on it), but for many of these the mutational history is uncertain: 87,406 have SPRTA placement support *<* 90%, and 53,365 *<* 50%. Of the remaining 1,436,089 genomes that are not mutationally informative (a terminal branch of length 0 separates them from the rest of the tree), 162,100 have SPRTA placement support *<* 90%, and 115,358 *<* 50%.

Phylogenetic uncertainty also affects many internal branches of the phylogenetic tree, highlighting substantial uncertainty in the ancestral mutational history, and not only in terminal branches: of 453,976 internal branches with inferred mutations, 59,523 have SPRTA support *<* 90%, and 29,641 *<* 50%. In terms of inferred mutations (which are assigned the same SPRTA score as the branch they are inferred to have occurred on), out of a total of 1,827,786 those with SPRTA support *<* 90% number 249,092 and there are 126,308 with SPRTA support *<* 50%.

One of the most striking examples of uncertainty in the SARS-CoV-2 mutational history is the evolution of the AY.4 Delta sub-lineage, one of the most abundant SARS-CoV-2 lineages, represented here by *>* 480, 000 genomes. Two mutations appear ancestral to most AY.4 genomes: T17040C and C21846T (Fig. 4, centre). After the appearance of mutation T17040C, the reversion C17040T appears to have occurred many times: we infer around 650 reversions. Position 17040 is inferred by MAPLE to have a substitution rate 31.9 times higher than average[21], mostly as a result of these reversions within AY.4. While inferred reversions are often due to reference biases in consensus genome calling methods, our consensus genomes were called with Viridian[22] which is not affected by this issue[21, 22]. Furthermore, we did not observe any issues with read data or substitution distribution along the tree that would suggest the presence of recurrent sequence errors at position 17040[21]. This suggests that 17040 is a genuinely hyper-mutable SARS-CoV-2 genome position, but only when in the mutated C nucleotide state, not the ancestral T. This is in line with the observation that mutation patterns in SARS-CoV-2 are highly position- and nucleotide-specific[23].

**Figure 4:**
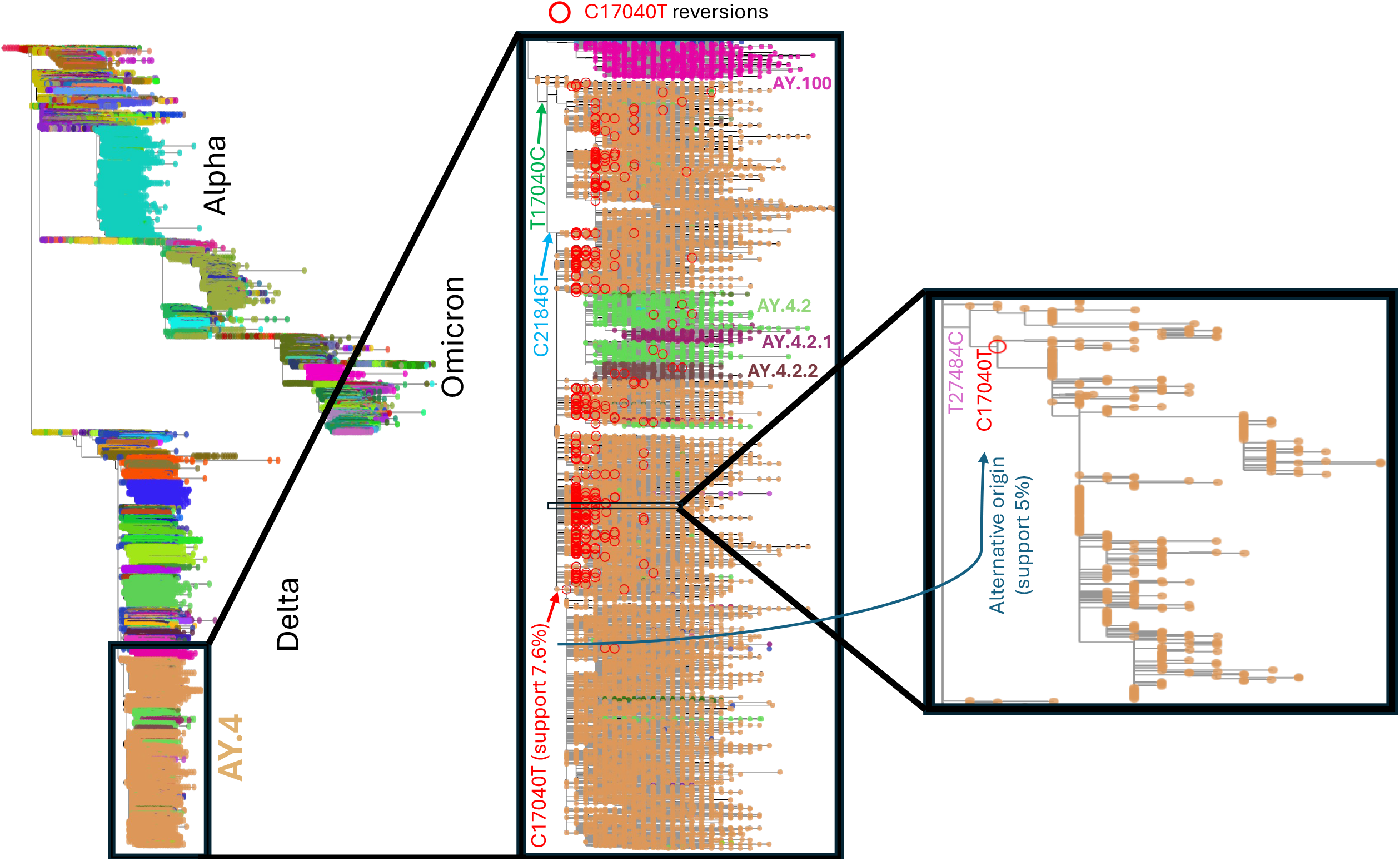
Uncertain evolutionary history of SARS-CoV-2 lineage AY.4. On the left is the global SARS-CoV-2 phylogenetic tree inferred by MAPLE and visualized in Taxonium[24]. Phylogenetic tips are coloured according to the Pango[11] lineage assigned by Pangolin[25] v4.3 (with pangolin-data v1.21) to the corresponding genomes. We also show the names of some of these lineages. In the centre we zoom-in on lineage AY.4, and show the locations of inferred T17040C (green arrow) and C21846T (blue arrow) mutations defining the inferred early evolution of the lineage. We highlight with red circles all *≈* 650 C17040T reversions within AY.4. One of them, shown with a red arrow, is ancestral to *>* 163, 000 genomes and has only 7.6% SPRTA support. On the right, we further zoom-in on the location of an alternative origin of this last sub-lineage (with corresponding SPR move represented by the dark blue arrow) that has 5% SPRTA support and entails re-placing the aforementioned sub-lineage as a direct descendant of the phylogenetic node with genome containing the T27484C and C17040T mutations.

The largest inferred sub-lineage within AY.4 descending from a C17040T reversion contains *>* 163, 000 genomes (Fig. 4, centre). However, the branch defining this lineage has only 7.6% SPRTA support. The reason is that, due to the hyper-mutability of C17040, there are many alternative plausible mutational histories within AY.4. The most likely alternative origin of this sub-lineage has SPRTA support of 5.0% and involves replacing the C17040T reversion defining this lineage with a C27484T reversion within a genomic background that already contains the C17040T reversion (Fig. 4, right). Although many equally parsimonious alternative mutation histories exist, this one is inferred by SPRTA to be the most likely alternative origin of this sub-lineage because the background mutation rate from C to T is very high in SARS-CoV-2, and because position 27484 also has an inferred substitution rate 20.8 times above average [21]. This example shows how SPRTA scores can not only highlight uncertain parts of an inferred phylogenetic tree and mutational history, but also identify and probabilistically assess alternative evolutionary origins of considered pathogen variants.

Regarding samples with uncertain placement, we show two examples (of many) in Figure 5. In the top example, uncertainty is caused by the sample having an incomplete genome sequence; in the bottom example, by the existence of two mutually plausible mutational histories. This shows how SPRTA can effectively identify the uncertainty related to the placement of rogue taxa, which by definition will have many possible placements but all with similarly low SPRTA support. Other branch support methods, in contrast, do not provide an evaluation of sample placement, as they only measure support for internal branches of the phylogenetic tree. Also, unlike for FB and most other cladistic branch support measures, rogue taxa mostly do not affect the SPRTA support scores of ancestral nodes, as their placement typically only affects the inference of mutation events on phylogenetic branches near the placement itself.

**Figure 5:**
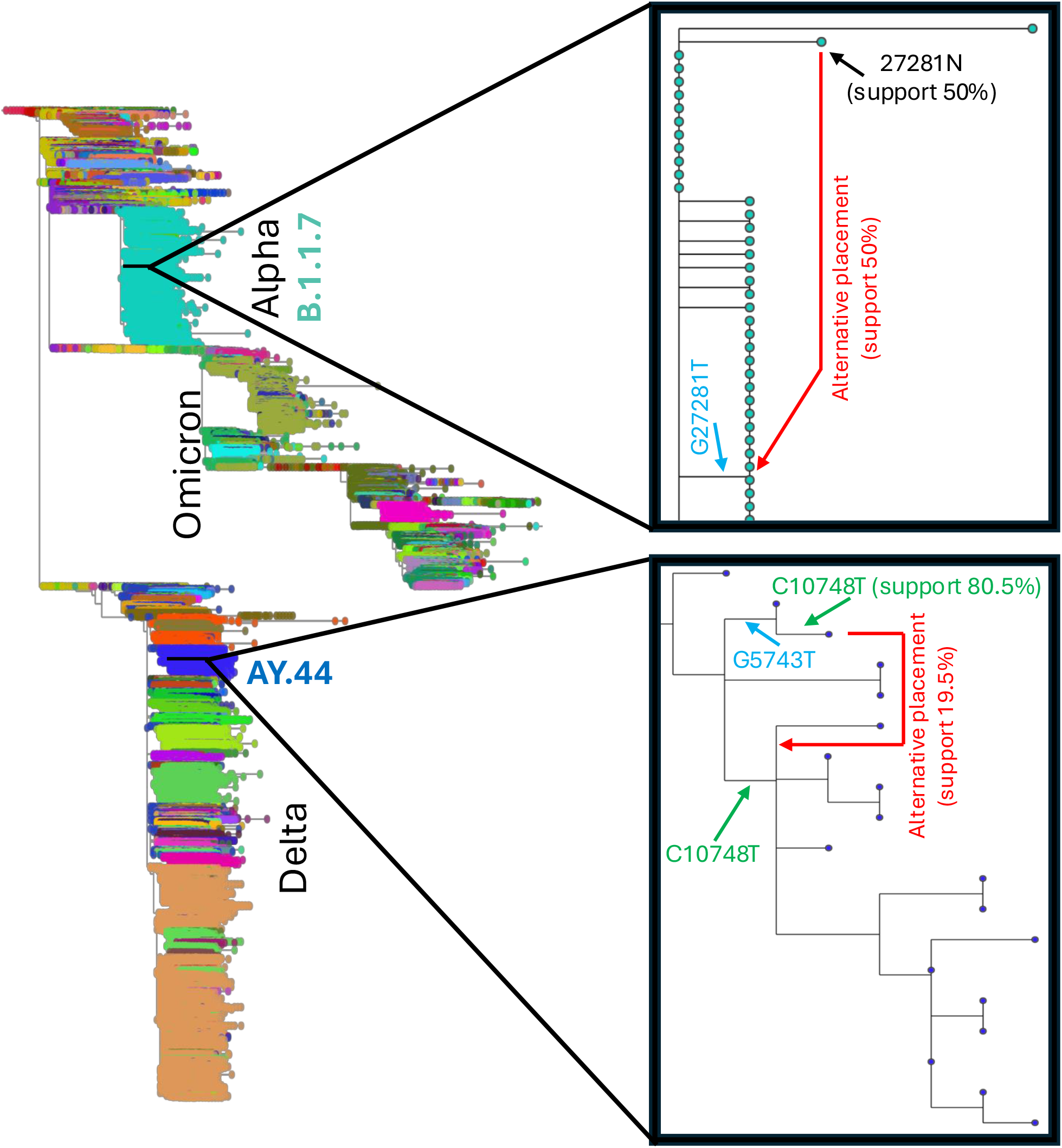
Sample placement uncertainty. To left is the global SARS-CoV-2 tree as in Figure 4. To right we zoom-in on two uncertain sample placements. At top, within lineage Alpha, the considered sample (marked by the black arrow) has no sequence information at genome position 27281 (ambiguity character N), the same position where a GT mutation occurs nearby in the phylogenetic tree (marked by the blue arrow). As such, this sample has 50% SPRTA support both at the current placement and at the other side of G27281T (marked by the red arrow). At bottom, within the AY.44 lineage, the placement of a sample has 80.5% SPRTA support, since the mutation C10748T implied by the current placement of the sample (upper green arrow) also occurs on a nearby branch (lower green arrow), permitting an alternative placement with 19.5% SPRTA support (red arrow). This alternative placement would require one fewer C10748T mutation on the tree but one additional G5743T mutation.

These are only some examples of all mutations and placements in the SARS-CoV-2 tree that are substantially uncertain (126,308 out of all 1,827,786 inferred mutations have SPRTA support *<* 50%.). Our full maximum likelihood tree annotated throughout with SPRTA support scores concisely represents the evolution of SARS-CoV-2 during the COVID-19 pandemic and presents the SPRTA probabilistic assessment of each sampled genome placement and each inferred mutation. It also offers a concise summary of plausible alternative placements and mutational histories. The annotated tree is available on Zenodo[26] together with the considered SARS-CoV-2 genome alignment, and can be visualized easily within Taxonium[24].

## 3 Discussion

With the increasing use of genomic epidemiology, pandemic-scale phylogenetics is set to become an essential tool for pandemic preparedness and epidemiology in general. New approaches such as MAPLE[18] and UShER[27] can be used to infer such massive phylogenetic trees, but there is inherently large uncertainty in these estimates due to complications including recurrent mutations and errors and incomplete genome sequences[21, 28, 29].

Traditional methods to quantify and represent this uncertainty cannot be used with pandemic-scale datasets. To overcome this problem, we present a new approach, SPRTA. In addition to addressing the limitations of existing methods in terms of computational demand, SPRTA also offers a new interpretation of branch support scores that is particularly useful in genomic epidemiology, replacing the cladistic focus of previous approaches.

Due to the excessive computational demand of applying Bayesian phylogenetic methods or Felsenstein’s bootstrap to large genome datasets, pandemic-scale phylogenies are usually taken at face value in downstream analyses such as inference of viral geographic spread[30], mutational patterns[23], lineage assignment[31], recombination[32], and variant fitness advantage[33]. Unchecked errors and uncertainty in phylogenetic trees can therefore propagate in these analyses and impact their accuracy and measures of uncertainty. Our approach allows us to efficiently distinguish between reliable and unreliable parts of an inferred phylogenetic tree, so that downstream analyses can focus on reliable phylogenetic signals or integrate over phylogenetic uncertainty.

While SPRTA is currently implemented in MAPLE, and is therefore optimized for applications to datasets with a low level of divergence, it is possible to implement the same approach within any phylogenetic inference tool, and so apply the same principles to any phylogenetic analysis.

SPRTA can also be used to create a phylogenetic network formed by a backbone maximum likelihood phylogenetic tree, extended to include additional branches with lower but substantial support. This way, SPRTA can be used to efficiently summarize vast numbers of possible phylogenetic trees (see, e.g., [34]). In the future, this might serve as a foundation for developing an efficient and complementary approach to Bayesian phylogenetics, helping to account for tree uncertainty in applications such as phylodynamics[35].

In conclusion, SPRTA not only addresses a fundamental outstanding problem in genomic epidemiology, but also offers a new paradigm in evolutionary biology for the interpretation and representation of phylogenetic information and uncertainty.

## 4 Methods

### 4.1 Definition of SPRTA and comparison to aBayes

Given an estimated phylogenetic tree *T* and data *D* in the form of a multiple sequence alignment, our aim is to assign confidence scores to branches *b* of *T*. We denote by *S*_*b*_ the subtree of a rooted tree *T* containing all descendants of *b*.

We take inspiration from the approximate Bayes (“ aBayes”) approach[16]. aBayes assigns to *b* a probability score *Pr*(*T* |*D*)_*b*_ based on the ratio of the likelihood *Pr*(*D*|*T*) of the estimated binary tree *T* vs. the likelihoods of the trees 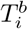 obtained using nearest neighbour interchange[17] (“ NNI”) moves centred around *b* (which comprise 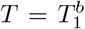 itself in addition to two tree topologies not containing *b*):

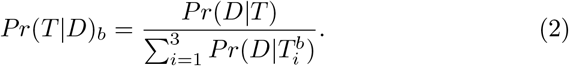

These NNI moves perform small changes to *T* near the location of branch *b*. This score is interpreted as an approximate Bayesian posterior score for *b*, where a flat tree prior is assumed, where instead of integrating over branch lengths (as typical in Bayesian phylogenetics) we define 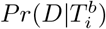 as the maximum likelihood score of topology 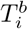 over all possible branch lengths, and where the only alternative topologies considered are those obtainable through a single NNI move starting from *T* [16]. *One of the appeals of aBayes is that it can score not only T*, but also the considered alternative topologies 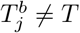:

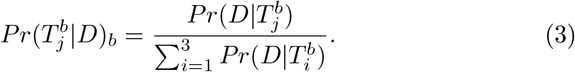

aBayes has a cladistic focus, and its support score for branch *b* is interpreted as the support for the clade containing all descendants of *b*. While much more computationally efficient than FB, aBayes can be too computationally demanding for large genomic epidemiological datasets if implemented within classical maximum likelihood phylogenetic methods (e.g. see Fig. 2). Also, aBayes is defined based on NNI moves, an approach insufficiently comprehensive for pandemic-scale data[18]: due to the existence of many phylogenetic topologies with similar likelihood[29], the only two alternative topologies obtainable through NNI moves and not containing the clade defined by branch *b*, 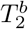 and 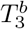, might represent only a very small subset of plausible alternative topologies not containing this clade.

Here, we address these limitations and define a new measure of branch support, SPRTA, that is particularly useful in the context of large-scale genomic epidemiology, but is also applicable more generally in phylogenetics. First, to address the problem of computational demand, we consider trees estimated using methods suitable for pandemic-scale datasets, such as MAPLE[18]. In the following, we assume that the likelihood of alternative tree topologies is also calculated with MAPLE or a similar method.

Within SPRTA, we define the support of branch *b* by considering a certain number *I*_*b*_ of possible alternative topologies 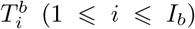, obtained by performing single subtree prune and regraft[17] (“ SPR”) moves that relocate *S*_*b*_ as a descendant of other parts of *T* (Fig. 1). Again, we assume for simplicity that 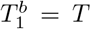 corresponds to the null SPR move. Compared to NNI moves, SPR moves are much more numerous, and can cause long-range changes to the tree; as such, SPR moves create a much more comprehensive set of alternative evolutionary histories than NNI moves. For a branch *b*, the number of possible SPR moves is linear in the number of sequences in the considered dataset, which means if we do not select which of these SPR moves we focus on, *I*_*b*_ can become too large, and the calculation of SPRTA scores too computationally demanding. However, to obtain an accurate evaluation we only need to consider topologies that have non-negligible likelihood score compared to *T* .

For this reason, we first perform an initial, approximate evaluation of alternative tree topologies obtained through SPR relocations of *S*_*b*_ using fixed branch lengths. From this, we only retain topologies with an initial log-likelihood score difference from *T* within a threshold corresponding approximately to one extra mutation event (the logarithm of the genome length, which, for SARS-CoV-2, is about 10.3). The likelihoods of all *I*_*b*_ topologies passing this threshold are then more deeply evaluated by optimizing branch lengths. More precisely, when we considering an alternative placement on branch *b*^*′*^ of sub-tree *S*_*b*_, let us call *n*^*′*^ the new node at the conjunction of *b*^*′*^ with *S*_*b*_. For this placement, the three branches whose length we optimize are the one connecting *n*^*′*^ with *S*_*b*_ (length *l*_1_), the one within *b*^*′*^ below *n*^*′*^ (length *l*_2_), and the one within *b*^*′*^ above *n*^*′*^ (length *l*_3_); see Supplementary Figure S2. 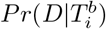 is defined as the maximum likelihood obtained optimizing these three branches, but leaving all other model parameters and branch lengths unaltered. This approximation of *Pr*(*D*|*T*^*b*^) is the same made by MAPLE, and is justified by the fact that, at the low levels of divergence typically considered in genomic epidemiology, changes in topology and branch lengths usually only affect nodes near those directly impacted by the changes[18].

Due to the initial filtering of plausible alternative topologies, *I*_*b*_ depends on branch *b*. These topologies are the same ones typically considered during the final stage of standard tree search in MAPLE.

SPRTA defines the support probability of *b* as

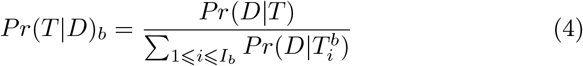

and we can similarly define 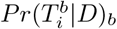 for alternative placements 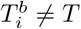 for any subtree *S*_*b*_ considered. Note that SPRTA typically evaluates a much larger number of alternative topologies compared to aBayes (Equation 2).

When evaluating different SPR placements, placements resulting in the same topology are equivalent, and so we only consider them once. In particular, we take into account multifurcations caused by branches of length 0, and consider placements into points of the tree with a null distance between them as equivalent (considering optimized placement-specific branch lengths as just described).

Because relative likelihood calculations of alternative topologies are performed as a standard component of tree inference in MAPLE, assessing SPRTA support probabilities adds negligible computational overhead.

### 4.2 Benchmarking of branch support methods

Branch support measures are often used to assess the expected accuracy of phylogenetic inference. However, how do we assess the accuracy of these measures of accuracy? We approach this problem using simulations, for which we have a ground truth against which to compared estimates. First, we simulate a tree and a set of genomes (Supplementary Fig. S3A); secondly, we estimate a tree from these simulated genomes (Supplementary Fig. S3B). Some branches of the inferred tree will be correct and some will be wrong: thirdly, we assess branch support measures according to their ability to give higher support scores to correctly estimated branches, and lower support scores to wrongly estimated branches (Supplementary Fig. S3C).

We benchmark branch support methods on simulated SARS-CoV-2 genome data (see Section 4.5.2) using an “ mutational” focus rather than a traditional “ cladistic” focus. We define phylogenetic correctness in terms of the genome evolutionary history implied by a phylogenetic tree. From each individual simulated dataset (see graphical example in Supplementary Fig. S3A), we first infer in MAPLE a maximum likelihood phylogenetic tree *T* and mutation events by marginal posterior mutation mapping[36] conditional on *T* (Supplementary Fig. S3B). Only mutation events inferred with *>* 0.5 probability by MAPLE are considered here. We define estimated mutation events as pairs (*G, m*) where *G* is a whole genome sequence and *m* = (*n*_1_, *p, n*_2_) is a single-nucleotide substitution at position *p* of *G* from nucleotide *n*_1_ (contained in *G* at position *p*) to nucleotide *n*_2_ (Supplementary Fig. S3C). An inferred mutation event (*G, m*) is considered correct if it is also present in the simulated tree, otherwise it is considered as an estimation error (Supplementary Fig. S3C).

Unlike cladistic methods, we identify an inferred phylogenetic branch *b* with the mutation events inferred to have occurred on *b*, not with the clade of taxa that descend from *b* in the inferred tree. As such, we consider the support score of *b* (estimated by SPRTA or any other method) as also the support score assigned to all the mutations inferred to have occurred on *b*. We consider a branch support measure as more accurate if in our simulations it assigns higher support to correctly estimated mutation events, and lower support to erroneously inferred ones.

### 4.3 Implementation and usage of SPRTA

We implemented SPRTA within MAPLE v0.6.8 (https://github.com/NicolaDM/MAPLE). While SPRTA can be run in MAPLE at the same time as tree inference, to aid comparability of computational performance with other approaches here we have considered its use to assess a pre-estimated input phylogenetic tree provided with the option “ *--inputTree*” . We used options “ *--numTopologyImprovements 0 --doNotImproveTopology*” to perform a shallow SPR search in MAPLE. We also used options “ *--model UNREST --rateVariation*” to use an UNREST model[37] with rate variation[21], and option “ *--estimateMAT*” to infer mutation events.

### 4.4 Other branch support methods

All other branch support measures considered here were calculated using IQ-TREE v2.1.3 with options “ *--seqtype DNA --seed 1 -m GTR+F+G4 –quiet -nt 1* “ . As with SPRTA, we always use the tree estimated by MAPLE as a starting tree (supplied via the option “ *-t* “) since on these datasets IQ-TREE will typically not converge to a tree with likelihood as high as MAPLE[18]. We used the following additional IQ-TREE options:

- “ *-B 1000* “ (1,000 bootstrap replicates) for UFBoot2[5].
- “ *--fast -b 100*” (100 bootstrap replicates and fast tree search) for Felsenstein’s bootstrap[1].
- “ *--fast -b 100 --tbe*” (100 bootstrap replicates and fast tree search) for TBE[6].
- “ *--fast --alrt 1000*” (1,000 bootstrap replicates and fast tree search) for aLRT-SH[15].
- “ *--fast --alrt 0*” (fast tree search) for aLRT[14].
- “ *--fast --abayes*” (fast tree search) for aBayes[16].
- “ *--fast --lbp 1000*” (1,000 bootstrap replicates and fast tree search) for LBP[13].

The number of bootstrap replicates has very limited impact on the computational demand of UFBoot2 and LBP, hence why these were set to 1,000.

### 4.5 SARS-CoV-2 genome datasets

#### 4.5.1 Viridian genome dataset

We applied SPRTA to a SARS-CoV-2 dataset containing 2,072,111 genomes collected up to February 2023. The consensus sequences of these genomes were consistently called with Viridian[22], a tool that prevents common reference biases in genomic regions of low sequencing coverage. Furthermore, we filtered out potentially contaminated samples, and masked alignment columns affected by recurrent sequence errors[21]. We estimated a phylogenetic tree using MAPLE v0.6.8 with an UNREST substitution model, rate variation, and deep SPR phylogenetic search. For a full description of data preparation and phylogenetic inference, see [21]. We ran SPRTA on this alignment and tree using MAPLE v0.6.9, with the options described in Section 4.3, and additionally with option *--supportFor0Branches* to evaluate the support of all genome placements, even those not involving mutations. The alignment, metadata, inferred tree, and SPRTA support scores are available on Zenodo [26].

#### 4.5.2 Simulated genomes

For benchmarking, we simulated SARS-CoV-2 genomes evolving along a known (“ true”) background phylogeny. The background tree we used was the publicly available 26 October 2021 global SARS-CoV-2 phylogenetic tree from http://hgdownload.soe.ucsc.edu/goldenPath/wuhCor1/UShER_SARS-CoV-2/[38] representing the evolutionary relationship of 2,250,054 SARS-CoV-2 genomes, as obtained using UShER[27].

We used phastSim v0.0.3[39] to simulate sequence evolution along this tree according to SARS-CoV-2 non-stationary neutral mutation rates[23], using the SARS-CoV-2 Wuhan-Hu-1 genome[40] as root sequence, and with gamma-distributed (*α* = 0.2) rate variation[41], similar to that estimated from real data[21].

## 5 Supplementary Figures

**Figure S1:**
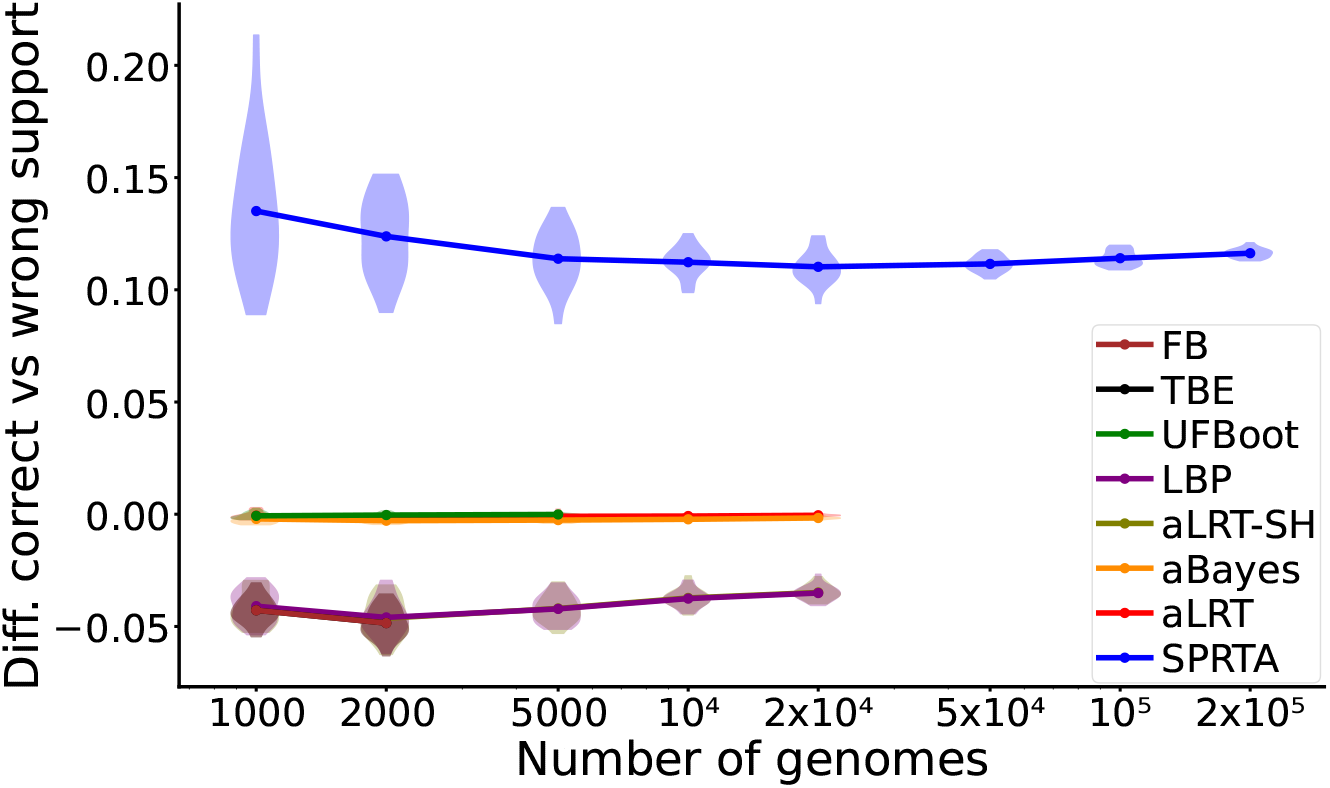
Difference between support scores of correct branches and wrong branches. On the Y axis we show the difference between the mean support of correctly inferred mutation events, and the mean support of wrongly inferred mutation events. All other details are as in Figure 3.

**Figure S2:**
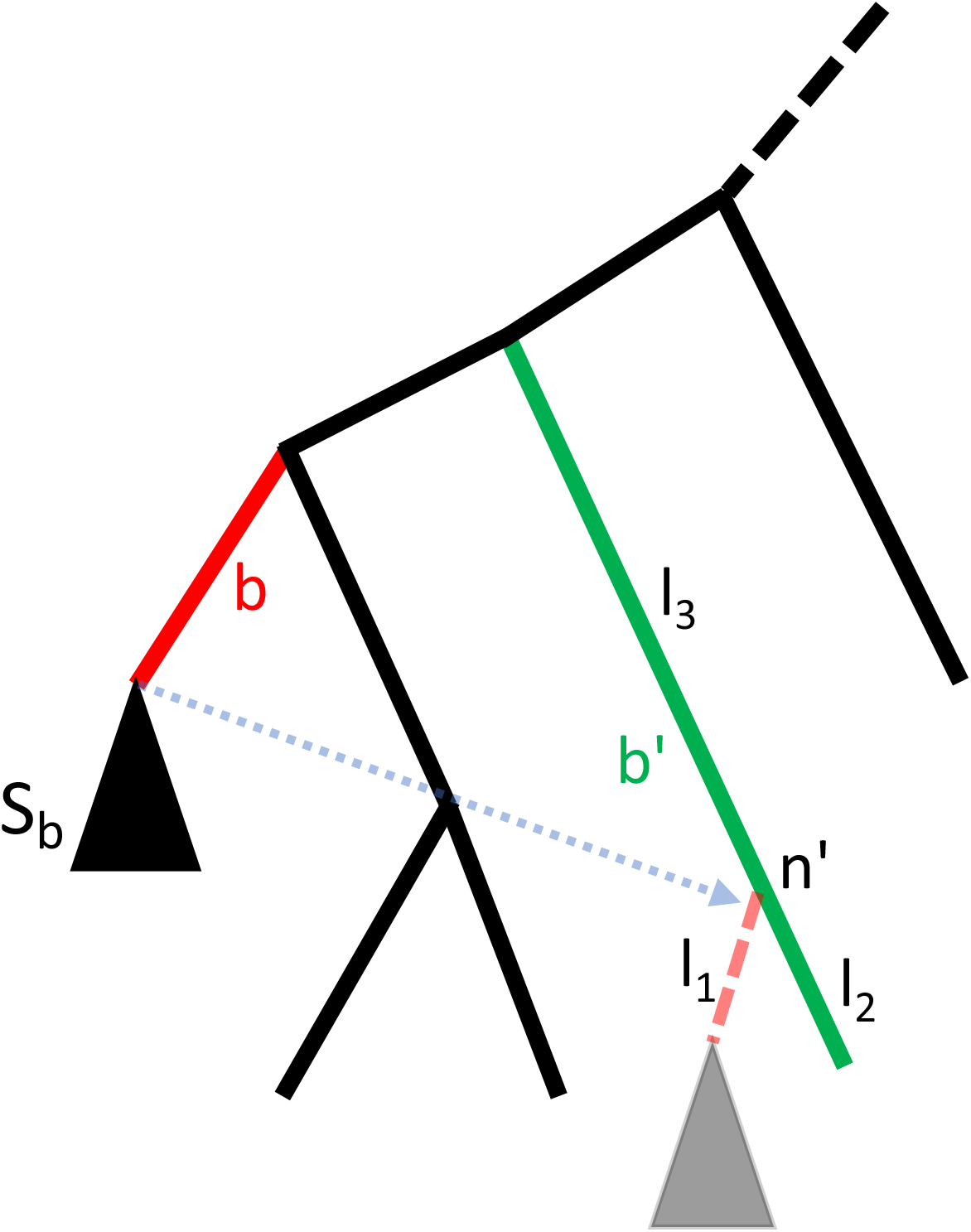
Graphical representation of branch length optimization for alternative topologies. Given a branch *b* (highlighted in red), we show an example re-placement (dotted shaded blue arrow) of its subtree *S*_*b*_ (black triangle). *n*^*′*^ is the node at which *S*_*b*_ is re-attached, within branch *b*^*′*^ (green). To evaluate this SPR move, we optimize the branch lengths *l*_1_, *l*_2_ and *l*_3_ in the figure, while the lengths of the rest of the branches in the tree are kept constant. Other details are as in Figure 1.

**Figure S3:**
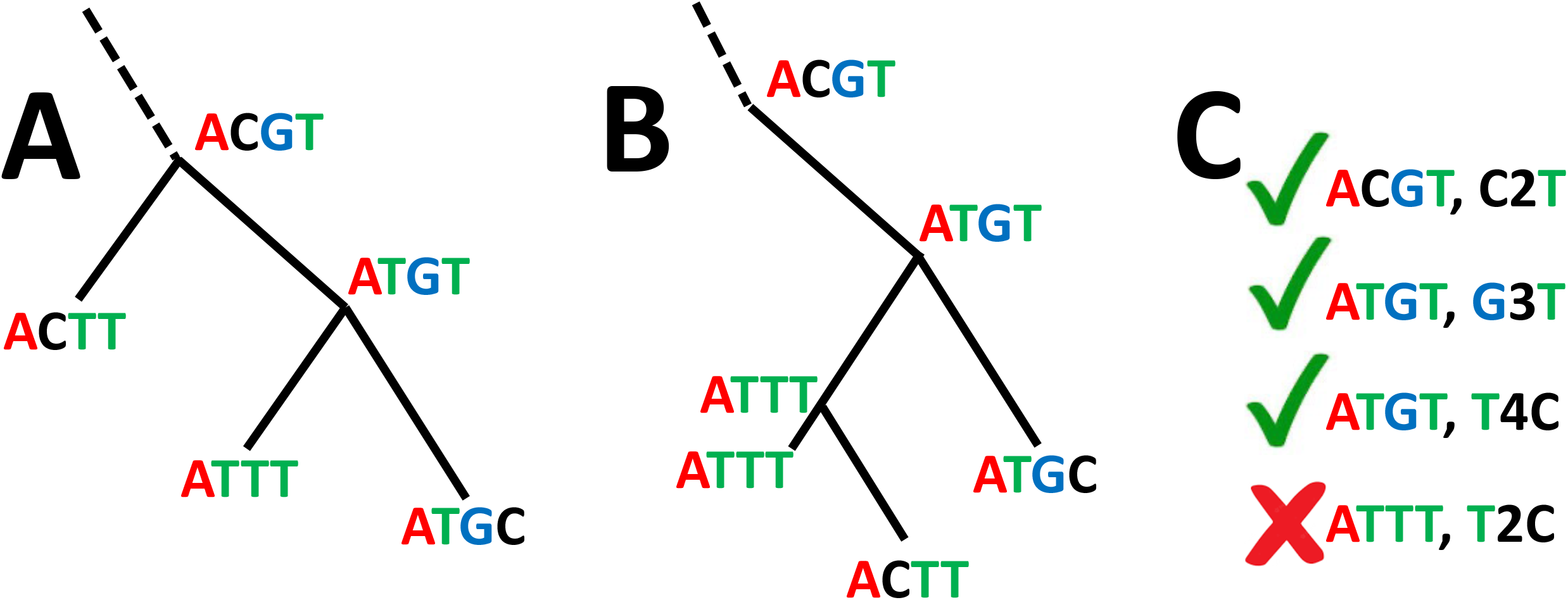
Graphical example of branch support assessment. To assess SPRTA, we consider support values given to correctly and wrongly inferred mutational events. Here we give a graphical example. **A** Part of an example true, simulated tree (the dashed branch represents the remainder of the tree) annotated with simulated ancestral genomes. For simplicity, we consider a genome of length 4. **B** Example tree estimated from genomes in **A**, annotated with inferred ancestral genomes. **C** Mutation events inferred in **B.**The first three mutation events are also present in **A**, and so are classified as correctly inferred. The final mutation event is instead classified as wrongly inferred. We evaluate our branch support method based on the score given to branches with correctly inferred mutations (where higher scores are considered better) vs. branches with wrongly inferred mutations (lower scores considered better).

## Notes

https://github.com/NicolaDM/MAPLE

### Competing Interest Statement

The authors have declared no competing interest.

